# Sex chromosome evolution in muscid flies

**DOI:** 10.1101/655845

**Authors:** Richard P. Meisel, Pia U. Olafson, Kiran Adhikari, Felix D. Guerrero, Kranti Konganti, Joshua B. Benoit

## Abstract

Sex chromosomes and sex determining genes can evolve fast, with the sex-linked chromosomes often differing between closely related species. A substantial body of population genetics theory has been developed and tested to explain the rapid evolution of sex chromosomes and sex determination. However, we do not know why the sex-linked chromosomes differ between some species pairs yet are relatively conserved in other taxa. Addressing this question will require comparing closely related taxa with conserved and divergent sex chromosomes and sex determination systems to identify biological features that could explain these rate differences. Cytological karyotypes suggest that muscid flies (e.g., house fly) and blow flies are such a taxonomic pair. The sex chromosomes appear to differ across muscid species, whereas they are highly conserved across blow flies. Despite the cytological evidence, we do not know the extent to which muscid sex chromosomes are independently derived along different evolutionary lineages. To address that question, we used genomic data to identify young sex chromosomes in two closely related muscid species, horn fly (*Haematobia irritans*) and stable fly (*Stomoxys calcitrans*). We provide evidence that the nascent sex chromosomes of horn fly and stable fly were derived independently from each other and from the young sex chromosomes of the closely related house fly (*Musca domestica*). We present three different scenarios that could have given rise to the sex chromosomes of horn fly and stable fly, and we describe how the scenarios could be distinguished. Distinguishing between these scenarios in future work could help to identify features of muscid genomes that promote sex chromosome divergence.

## Introduction

In species where sex is determined by genetic differences between males and females, sex determining loci can reside on sex chromosomes, such as the male-limited Y chromosome in mammals that carries a male-determining gene (Swain and Lovell-Badge, 1999). When the X and Y (or Z and W) chromosomes are highly differentiated, the Y (or W) chromosome contains only a handful of genes with male-specific functions (Bachtrog, 2013; Charlesworth *et al.*, 2005). X (or Z) chromosomes, on the other hand, typically resemble autosomes in gene density, with some differences in the types of genes found on the X and autosomes (Meisel *et al.*, 2012; Vicoso and Charlesworth, 2006). Other sex chromosome pairs are homomorphic, with little sequence differentiation between the X and Y (or Z and W) chromosomes (Wright *et al.*, 2016). Sex determining genes and sex chromosomes often differ across species because of evolutionary turnover as species diverge.

Sex determining genes and sex chromosomes often differ across species (Bachtrog *et al.*, 2014; Beukeboom and Perrin, 2014), predominantly as a result of two general processes. First, when a new sex determining locus arises on an autosome, it can convert the autosome into a “proto-sex-chromosome”, and the ancestral sex chromosome can revert to an autosome (Carvalho and Clark, 2005; Larracuente *et al.*, 2010; van Doorn, 2014). Second, autosomes can fuse with X, Y, Z, or W chromosomes to create “neo-sex-chromosomes” (Pennell *et al.*, 2015; Vicoso and Bachtrog, 2015). A special case of chromosomal fusions are reciprocal translocations, in which an autosomal region is translocated to a sex chromosome and *vice versa* (e.g., Toups *et al.*, 2019). Population genetics theory suggests that sex-specific selection pressures (including sexual antagonism) are important contributors to the evolution of sex determination pathways, evolutionary turnover in sex chromosomes, and the fixation of neo-sex chromosomes (Charlesworth and Charlesworth, 1980; Rice, 1986; van Doorn and Kirkpatrick, 2007, 2010). This theory has been tested in many plants and animals, and those tests have generally supported the hypothesis that sex-specific selection is important for the evolution of sex chromosomes and sex determination (e.g., Qiu *et al.*, 2013; Roberts *et al.*, 2009; Wang *et al.*, 2012; Zhou and Bachtrog, 2012).

In some taxa, the sex-linked chromosomes or sex determining genes differ across many species, whereas in other taxa the same X and Y (or Z and W) chromosomes are conserved across (nearly) all species (Beukeboom and Perrin, 2014; Blackmon and Demuth, 2014). Despite the well constructed theory explaining how sex chromosomes and sex determination evolve, and the empirical work supporting that theory, we know very little about why the rates of evolution differ across taxa (Abbott *et al.*, 2017; Bachtrog *et al.*, 2014). Contrasting closely related taxa with conserved and divergent sex chromosomes could allow for the identification of biological factors that affect sex chromosome divergence.

Brachyceran flies (i.e., higher Diptera) are well-suited models for identifying factors that promote or inhibit sex chromosome evolution because they have variable rates of sex chromosome divergence across lineages (Vicoso and Bachtrog, 2015). The karyotype of the most recent common ancestor (MRCA) of Brachycera consists of five gene-rich autosomes (Muller elements A–E), a gene-poor heterochromatic X chromosome (Muller element F), and a Y chromosome that carries a male-determining locus (**Fig 1**) (Boyes and Van Brink, 1965; Foster *et al.*, 1981; Vicoso and Bachtrog, 2013; Weller and Foster, 1993). Elements A–E correspond to the five gene-rich chromosome arms of *Drosophila melanogaster* (X, 2L, 2R, 3L, and 3R), and element F is homologous to the gene-poor *D. melanogaster* dot chromosome, i.e., chromosome 4 (Muller, 1940). Element F has been conserved as the X chromosome for ∼175 million years (My) in some phylogenetic lineages within flies, while new sex chromosomes have arisen along other lineages (Baker and Wilkinson, 2010; Vicoso and Bachtrog, 2013, 2015; Wiegmann *et al.*, 2011).

**FIG. 1.**
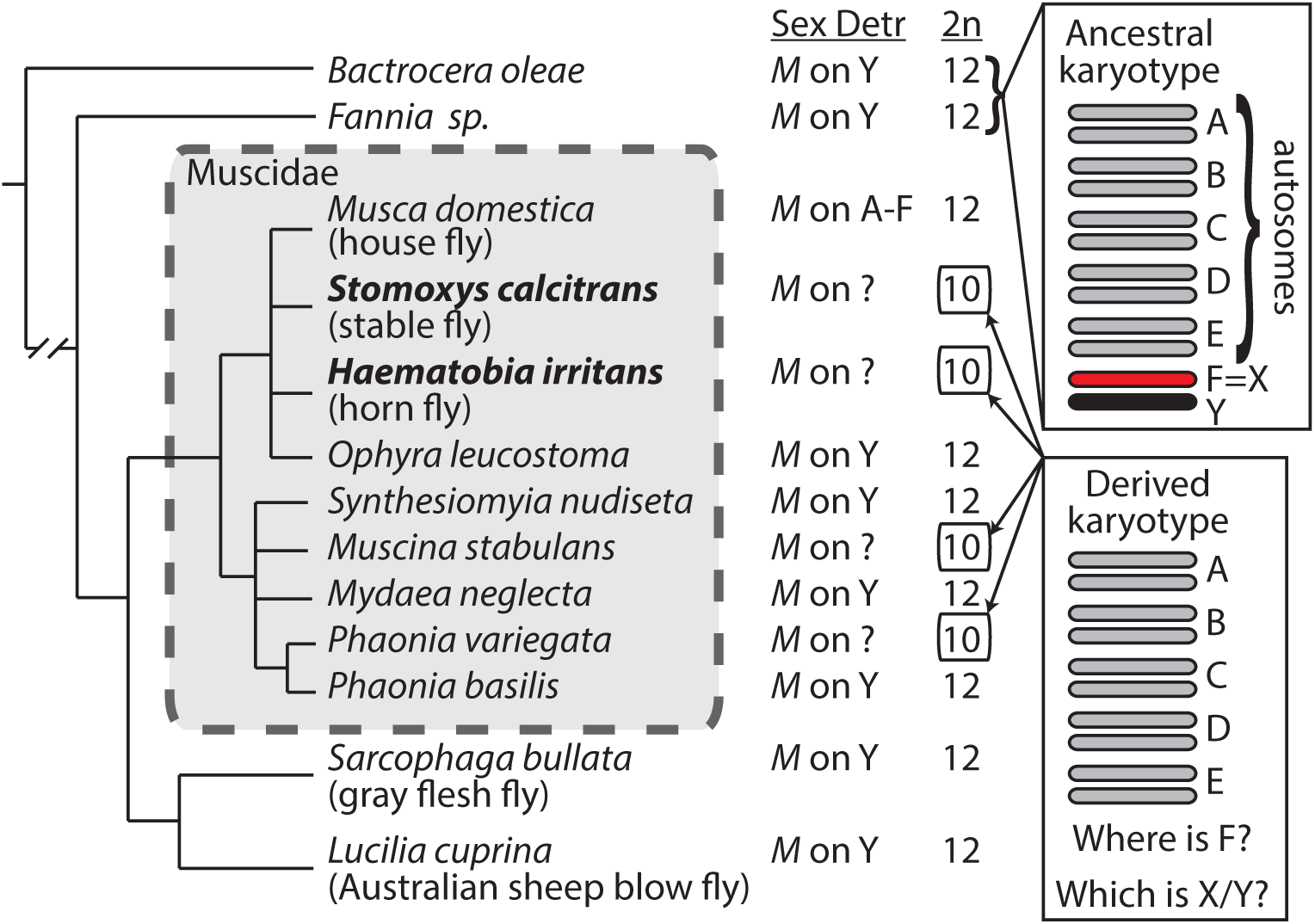
Cryptic sex chromosomes in Muscidae. Phylogenetic relationships and karyotypes of muscid flies and their relatives (Boyes and Van Brink, 1965; Boyes *et al.*, 1964; Schnell e Schuehli *et al.*, 2007; Vicoso and Bachtrog, 2015). The inferred mechanism of sex determination (Sex Detr) and the diploid chromosome number (2n) are listed for each species. “*M*” refers to a generic male-determining locus.

Within Brachycera, the sister families of Muscidae (house flies and their allies), Calliphoridae (blow flies), and Sarcophagidae (flesh flies) could be especially informative for comparative studies because they appear to have family-specific rates of sex chromosome evolution (**Fig 1**). These three families diverged from their common ancestor ∼50 My ago (Wiegmann *et al.*, 2011). Nearly all blow flies and flesh flies have the ancestral fly karyotype, with five autosomes, a heterochromatic X, and a male-determining locus on a Y chromosome (Boyes, 1961; Boyes and Van Brink, 1965; Scott *et al.*, 2014b; Vicoso and Bachtrog, 2015). The only exceptions are sex determination by maternal genotype in the blow fly *Chrysomya rufifacies* (Ullerich, 1963), and an expanded karyotype of 19–20 chromosomes in the flesh fly *Agria* (*Pseudosarcophaga*) *affinis* (Boyes, 1953). The X chromosomes of the Australian sheep blow fly (*Lucilia cuprina*) and gray flesh fly (*Sarcophaga bullata*) both correspond to element F (Vicoso and Bachtrog, 2013, 2015), suggesting that element F is the ancestral X of these families. In addition, the Y chromosomes of *L. cuprina* and *S. bullata* are extremely differentiated from their homologous X chromosomes, suggesting that they have existed as an X-Y pair for many millions of years (Vicoso and Bachtrog, 2013, 2015). Furthermore, the haploid X chromosome in *L. cuprina* males is up-regulated (i.e., dosage compensated) by an RNA-binding protein that is homologous to a *Drosophila* protein that binds nearly exclusively to element F (Davis *et al.*, 2018; Linger *et al.*, 2015). As expected because of the genetic differentiation between the *L. cuprina* X and Y, loss of function mutations in the *L. cuprina* gene encoding the dosage compensation protein are lethal specifically in males (Davis *et al.*, 2018).

In contrast to flesh flies and blow flies, multiple lineages in the family Muscidae seem to have evolved new sex chromosomes in the <40 My since the common ancestor of the family (Ding *et al.*, 2015). The iconic example of sex chromosome evolution in Muscidae is the house fly (*Musca domestica*), which has a well-characterized polygenic sex determination system (Hamm *et al.*, 2015). House fly appears to have the ancestral brachyceran karyotype (i.e., 5 pairs of euchromatic chromosomes and a heterochromatic sex chromosome pair), but the X and Y chromosomes are not differentiated (Meisel *et al.*, 2017). This is because the house fly male-determining locus (*Mdmd*) is a recently derived duplicated gene that can be found on at least 5 of the 6 chromosomes (Sharma *et al.*, 2017). The invasion and spread of this new male-determiner in the house fly genome since the divergence with closely related species is such that every house fly chromosome can be an undifferentiated proto-sex-chromosome pair.

In addition to house fly, other Muscidae have derived karyotypes that were evidenced by cytological examination (**Fig 1**). For example, the heterochromatic element F is missing from the karyotypes of stable fly (*Stomoxys calcitrans*), horn fly (*Haematobia irritans*), and some other muscids (Avancini and Weinzierl, 1994; Boyes *et al.*, 1964; Joslyn *et al.*, 1979; LaChance, 1964; Parise-Maltempi and Avancini, 2007), possibly because it fused to another element. Stable fly and horn fly both have genetic sex determination with a dominant male-determining locus (McDonald and Schmidt, 1987; Willis *et al.*, 1981), but the specific identities of the X and Y chromosomes remains unresolved. We hypothesize that species with these derived karyotypes have cryptic sex chromosomes that arose when an ancestral sex chromosome (element F and/or the Y chromosome) recently fused to one of the other five chromosomes. Here, we describe the identification of the cryptic sex chromosomes in stable fly and horn fly using genomic and transcriptomic sequence data. We also provide evidence that the stable fly and horn fly sex chromosomes are young and of independent origins. These results demonstrate that muscid flies are a good model system for studying the factors that permit rapid evolution of sex chromosomes.

## Methods

### Assigning scaffolds to Muller elements

We used a homology-based approach that we had previously developed in house fly to map stable fly and horn fly scaffolds to Muller elements (Meisel and Scott, 2018; Meisel *et al.*, 2015). This approach works because Muller element gene content (synteny) is conserved across Brachycera (Foster *et al.*, 1981; Sved *et al.*, 2016; Vicoso and Bachtrog, 2013; Weller and Foster, 1993). For stable fly, we selected OrthoGroups from the OrthoDB annotation that contain a single *D. melanogaster* gene and a single *S. calcitrans* gene (Kriventseva *et al.*, 2018; Olafson *et al.*, 2019). For horn fly, we obtained annotated genes from the initial analysis of the genome, and we extracted the inferred *D. melanogaster* homologs for each gene (Konganti *et al.*, 2018). We assigned the stable fly and horn fly genes to the same Muller element as their *D. melanogaster* homologs. Each of these genes is part of a chromosomal scaffold. We used a majority-rules approach to assign those scaffolds to Muller elements if >50% of the genes on a scaffold are assigned to the same Muller element (Supplementary tables S1 and S2). This allowed us to assign 94.6% (1,482/1,566) of stable fly scaffolds and 97.5% (4,778/4,889) of horn fly scaffolds containing annotated genes to Muller elements. All genes on a scaffold are then assigned to the Muller element of that scaffold regardless of the Muller element designation of their annotated ortholog. Assigning genes to Muller elements based on their scaffold should control for individual genes that are positionally relocated between elements across flies (Baker and Wilkinson, 2010; Bhutkar *et al.*, 2007).

### Variant calling

Our approach to identifying sex chromosomes involves testing for Muller elements with increased heterozygosity in males, which is the expectation for young, undifferentiated X-Y chromosome pairs (Meisel *et al.*, 2017; Vicoso and Bachtrog, 2015). To those ends, we used the Genome Analysis Toolkit (GATK) version 3.4-0 to identify heterozygous single nucleotide polymorphisms (SNPs) in stable fly and horn fly genomes and transcriptomes, following the GATK best practices (DePristo *et al.*, 2011; McKenna *et al.*, 2010; Van der Auwera *et al.*, 2013). The Illumina sequencing reads used to assemble the stable fly genome were generated from DNA extracted from males of a strain that had been inbred for 7 generations (Olafson *et al.*, 2019). These data allow us to identify nascent sex chromosomes based on elevated male heterozygosity. The same cannot be done for horn fly because the sequences used to assemble the genome came from DNA isolated from mixed pools of males and females (Konganti *et al.*, 2018).

To quantify heterozygosity in stable fly males, we first mapped the sequencing reads to the assembled genomic scaffolds using the MEM algorithm implemented in BWA with the default parameters (Li and Durbin, 2009). Next, we used Picard Tools version 1.133 to identify and remove duplicate reads, and we realigned indels with GATK. Then, we performed näive variant calling using the GATK HaplotypeCaller with a phred-scaled confidence threshold of 20, and we selected the highest confidence SNPs from that first-pass (QD < 2.0, MQ < 40, FS > 60, SOR > 4, MQRankSum < −12.5, ReadPosRankSum < −8). We used those high quality variants to perform base recalibration, we re-input those recalibrated bases into another round of variant calling, and we extracted the highest quality variants. We repeated the process so that we had performed three rounds of recalibration, which was sufficient for convergence of variant calls. We applied GenotypeGVCFs to the variant calls from all of the Illumina libraries for joint genotyping. We then used the GATK HaplotypeCaller to genotype all of the variable sites (phred-scaled confidence > 20), and we selected only the high quality variants (FS > 30 and QD < 2).

Comparing male and female heterozygosity, rather than only analyzing male heterozygosity, may be a better way to identify elevated heterozygosity in males. There are no genomic sequences available from female stable fly or from single-sex horn fly to use in such an analysis. However, Illumina RNA-seq data were collected from female and male tissues separately for both species. From stable fly, RNA-seq libraries were sequenced from female and male whole adults (SRX229930 and SRX229931), reproductive tissues (SRX995859 and SRX995857), and heads (SRX995858 and SRX995860). RNA from whole adult stable flies was extracted from the same inbred strain that supplied the DNA for the genome assembly, and RNA from reproductive tissues and head was extracted from the stable flies in the lab colony from which the inbred line was derived. From horn fly, RNA-seq libraries were sequenced from ovary (SRX3340090) and testis (SRX3340086). Horn flies were sampled from the USDA-ARS Knipling-Bushland U.S. Livestock Insects Research Laboratory strain, which has been maintained since 1961 (Konganti *et al.*, 2018). One RNA-seq library was sequenced for each tissue from each species (Konganti *et al.*, 2018; Olafson *et al.*, 2019).

We used a modified GATK pipeline to identify SNPs in stable fly and horn fly RNA-seq data (Meisel *et al.*, 2017). First, RNA-seq reads were aligned to the reference genomes of the appropriate species using STAR version 2.4.0.1 (Dobin *et al.*, 2013). We used the aligned reads to create a new reference genome index from the inferred spliced junctions in the first alignment, and then we performed a second alignment with the new reference. We next marked duplicate reads and used SplitNCigarReads to reassign mapping qualities to 60 with the ReassignOneMappingQuality read filter for alignments with a mapping quality of 255. Indels were realigned, and three rounds of variant calling and base recalibration were performed as described above for the stable fly genomic sequencing reads. We applied GenotypeGVCFs to the variant calls from all tissues for joint genotyping of males and females from each species. Finally, we used the same filtering parameters that we applied to the stable fly genomic sequencing reads to extract high-quality SNPs from our variant calls.

Once we had identified sites in the sequence data that differ from the reference genome, we extracted heterozygous sites across the genome within annotated genes. We used those data to calculate the number of heterozygous SNPs per Mb within each annotated gene (Supplementary tables S3–S5). Genes with no heterozygous sites were omitted from the results we present, but we obtain the same general patterns when these genes are included. For each gene, we calculated the fraction of all heterozygous sites in either sex that are heterozygous in males. This value ranges from 0 (if heterozygous sites are only observed in females) to 1 (if heterozygous sites are only observed in males), with 0.5 indicating equal heterozygosity in males and females.

### Gene expression analysis

We aligned the same RNA-seq data described above to the annotated transcripts in the stable fly and horn fly genomes and calculated transcripts per million reads (TPM) for each transcript using kallisto version 0.44.0 (Bray *et al.*, 2016). We also used kallisto to align 454 GS FLX reads from whole adult female (SRR003191) and male (SRR003191) horn flies to the horn fly reference transcripts (Konganti *et al.*, 2018). In addition, we used the same approach to align Illumina RNA-seq reads from whole adult male (SRX208993 and SRX208994) and female (SRX208996 and SRX208997) house flies to the house fly reference genome (Scott *et al.*, 2014a). We then summed TPM for all transcripts from each gene for each sample type (e.g., stable fly female head) to obtain a gene-level estimate of expression in each sample (Supplementary tables S6–S11). In house fly, where there are two RNA-seq libraries for each sex, we calculated the mean TPM for each gene across both libraries. Using these data, we calculated the log2-fold male:female expression level 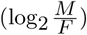 of each gene for each tissue type. In our analysis, we only considered genes with TPM > 0 in both males and females in a particular tissue and TPM > 1 in at least one sex.

## Results

### The stable fly sex chromosomes consist of elements D and F

We hypothesize that stable fly has young, cryptic sex chromosomes (**Fig 1**). Nascent sex chromosomes can be identified based on elevated heterozygosity in the heterogametic sex (i.e., XY males) because the X and Y have begun to differentiate in the sequences of genes, but the Y still retains most of the genes in common with the X chromosome (Meisel *et al.*, 2017; Vicoso and Bachtrog, 2015). The stable fly genome was sequenced from male DNA (Olafson *et al.*, 2019), allowing us to identify the sex-linked element(s) by testing for elevated heterozygosity in the genome sequencing reads. To those ends, we first assigned most of the genes in the stable fly genome to Muller elements using homology relationships with *D. melanogaster* (Supplementary table S1). Next, we identified heterozygous SNPs in the sequencing reads generated from stable fly males (Supplementary table S3). Because multiple males were sampled for genome sequencing, our approach will capture two different types of variable sites: fixed differences between the X and Y chromosomes, as well as polymorphisms on the X, Y, and autosomes that segregate in the lab strain that was sampled. We expect the sex chromosomes to have elevated heterozygosity sites, whereas the autosomes will only have the latter. We performed all of our analyses on heterozygous variants that we identified within annotated genes.

We found that stable fly genes assigned to Muller element D have more heterozygous SNPs in males than genes on scaffolds mapped to the other four major elements (*P* =10^−138^ in a Mann-Whitney test comparing element D genes with genes on elements A, B, C, and E; **Fig 2A**). We performed a similar analysis comparing each of the other four major elements against all others, and we did not detect any other elements with elevated heterozygosity. This suggests that element D is part of the stable fly X and Y chromosomes.

**FIG. 2.**
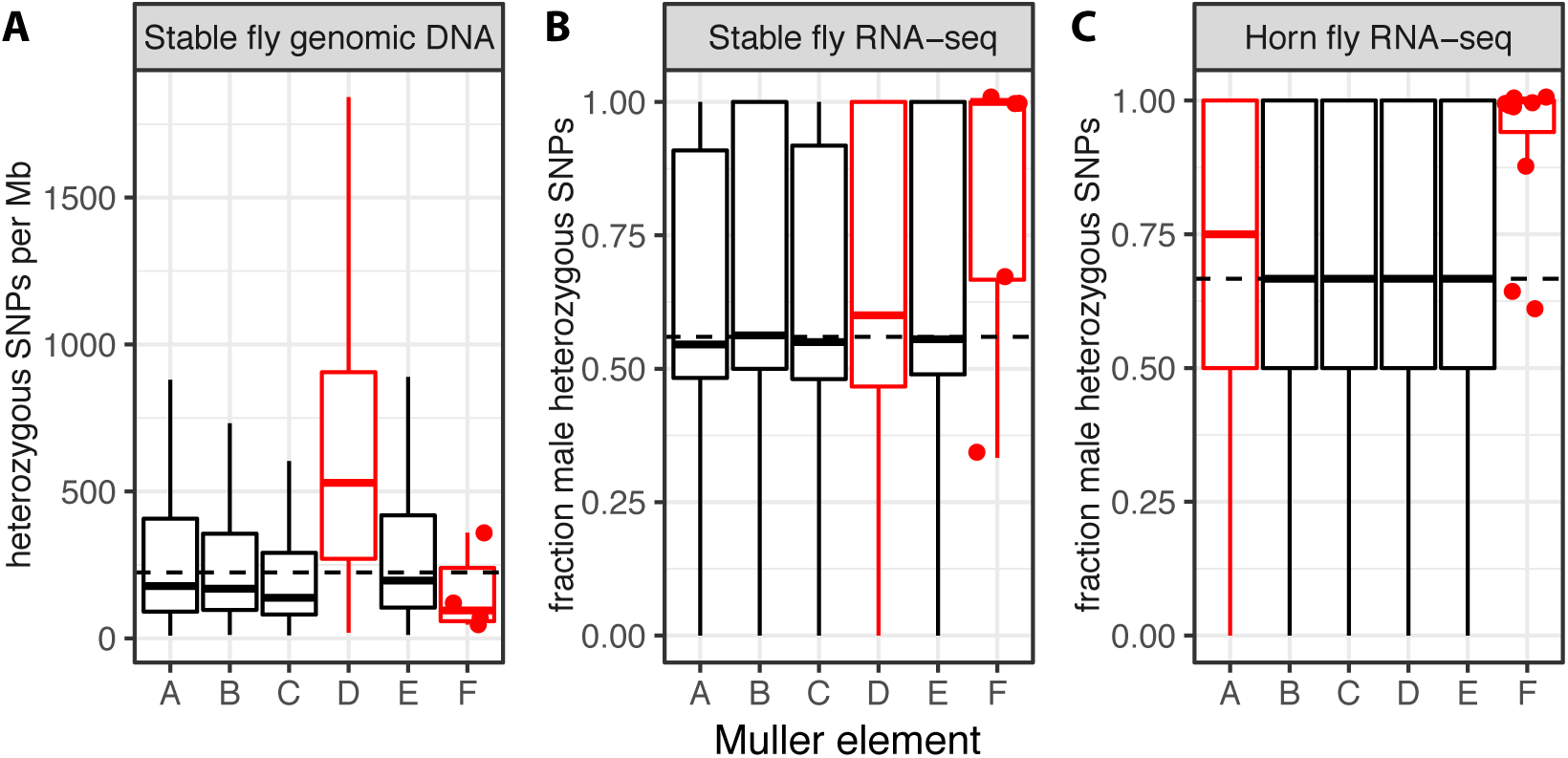
Identifying stable fly and horn fly sex chromosomes. **A.** Boxplots show the number of heterozygous SNPs per megabase (Mb) identified using genomic sequencing reads from males within annotated stable fly genes mapped to each of the Muller elements. **B–C.** Boxplots show the fraction of heterozygous SNPs in males relative to females within annotated stable fly and horn fly genes mapped to each of the Muller elements, using RNA-seq data. Each data point used to generate the boxplots corresponds to an individual gene. Dots indicate the values for element F genes. Dashed lines indicate the genome-wide average for all genes. Outliers were omitted from all plots. Inferred sex-linked elements are drawn in red.

We may have also expected elevated male heterozygosity on element F because it is the ancestral brachyceran X chromosome, but element F genes do not have an excess of heterozygous SNPs (**Fig 2A**). However, element F has reduced variation along most of the length in *Drosophila*, likely because a lack of recombination enhances the diversity-reducing effects of selective sweeps and background selection (Arguello *et al.*, 2010; Berry *et al.*, 1991; Hilton *et al.*, 1994; Jensen *et al.*, 2002; Wang *et al.*, 2002, 2004). Therefore, the low male heterozygosity of element F genes in stable fly could be explained if they also experience reduced recombination. A key limitation of this analysis is that we only examined heterozygosity in males, and our prediction is that because they will contain both types of variable there will be elevated male heterozygosity when compared to females (Vicoso and Bachtrog, 2015).

To compare male and female heterozygosity, we used available RNA-seq data to identify heterozygous SNPs separately in each sex (Olafson *et al.*, 2019). We then calculated relative male heterozygosity as the fraction of all SNPs in each gene that are heterozygous in males (Supplementary table S4). Relative male heterozygosity ranges from 0, if all heterozygous SNPs are in females, to 1, if all heterozygous SNPs are in males (Meisel *et al.*, 2017). As in our analysis of absolute male heterozygosity, we expect the sex chromosomes to have elevated relative male heterozygosity because sex-linked genes will harbor fixed differences between the X and Y chromosomes, in addition to segregating polymorphisms. Consistent with our analysis of absolute heterozygosity, there is elevated relative male heterozygosity in genes on element D (*P* =0.047 in a Mann-Whitney test comparing element D to the other four major elements; **Fig 2B**). None of the other major elements (A, B, C, or E) have elevated relative male heterozygosity. It is also curious that male heterozygosity is, on average, higher than female heterozygosity across the entire genome. We do not have an explanation for this. We cannot examine heterozygosity along the full length of any individual Muller elements because there is not a chromosome-scale assembly of the stable fly genome, and the order of the assembled scaffolds along each chromosome has not yet been determined.

Other comparisons of male and female heterozygosity support the sex-linkage of element D in stable fly. For example, there is a higher fraction of element D genes that only have heterozygous SNPs in males and not in females (518*/*1850 = 28.0%) when compared to genes on the other major elements (2168*/*8625 = 25.1%; *z*=2.56, *P* =0.005). Moreover, when we limit our analysis to only those heterozygous SNPs identified in both the genomic DNA and RNA-seq data, we find that an excess of element D genes have at least one heterozygous SNP (507*/*2437) when compared to genes on the other four major elements (503*/*11226; *P*<10^−15^ in Fisher’s exact test).

We also detect evidence for elevated relative male heterozygosity on stable fly element F (the ancestral X chromosome). Only 10 annotated stable fly genes are assigned to element F, of which five have heterozygous SNPs. Four of those five genes have more heterozygous SNPs in males than females. It is unlikely that *≥*4*/*5 of element F genes would have >50% male heterozygous SNPs if heterozygosity were equal in males and female (*z*=1.34, *P* =0.09). In addition, three of the five element F genes only have heterozygous SNPs in males, which is more than the frequency of autosomal genes (25.1%) that only have male heterozygous SNPs (*z*=1.80, *P* =0.04). These results support the hypothesis that the stable fly X and Y chromosomes are young and minimally differentiated. They also suggest that the X chromosome consists of elements D and F, as does the Y chromosome.

The genome assembly of stable fly provides a third line of support that element D has higher male heterozygosity and is therefore part of the X and Y chromosomes. We expect heterozygosity to interfere with genome assembly, leading to smaller contigs and scaffolds (Kajitani *et al.*, 2014; Pryszcz and Gabaldón, 2016; Vinson *et al.*, 2005). In the case of XY males with nascent sex chromosomes, this assembly fragmentation should be greater on sex-chromosome-derived scaffolds than autosomal scaffolds. More scaffolds in the stable fly assembly are assigned to element D than any of the other elements, even though element D does not have more genes or a larger inferred length than the other elements (**Fig 3A-C**). This suggests that the assembly of stable fly element D is more fragmented than the other four major elements. Consistent with this prediction, scaffolds assigned to stable fly element D are shorter (*P* =10^−15^ in a Mann-Whitney test) and have fewer genes (*P* =10^−15^ in a Mann-Whitney test) than scaffolds assigned to the other four major chromosomes (**Fig 3D-E**).

**FIG. 3.**
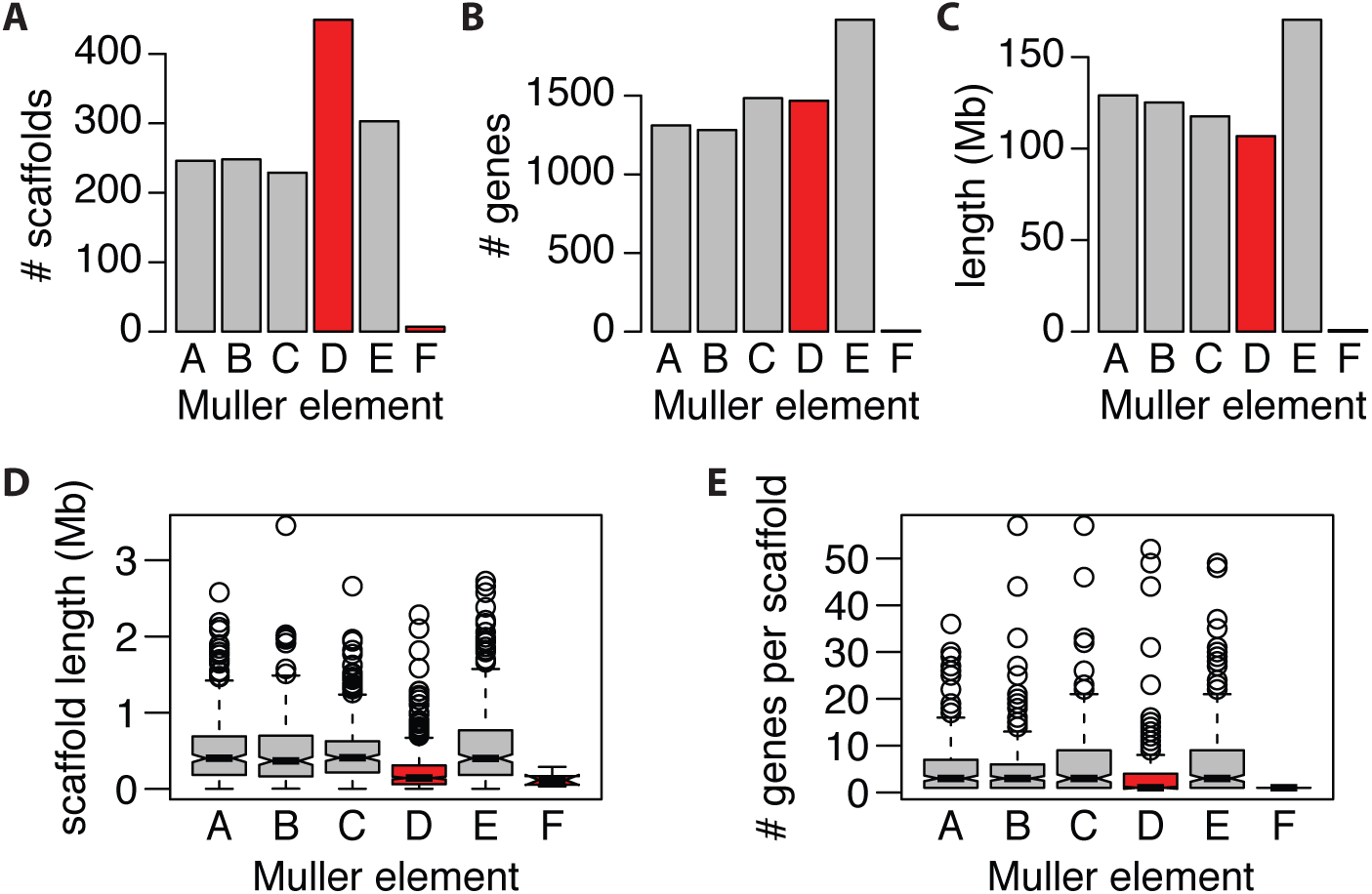
Fragmented assembly of the stable fly sex chromosome. **A.** The number of genomic scaffolds and **B.** the number of genes assigned to each stable fly Muller element. **C.** The composite length of all scaffolds assigned to each stable fly Muller element. **D.** The distributions of scaffold lengths and **E.** the distributions of genes per scaffold for scaffolds assigned to each stable fly Muller element are shown with boxplots. Inferred sex-linked elements are highlighted in red.

Therefore, there are three lines of evidence that are all consistent with elevated heterozygosity on element D in stable fly males, providing a congruent picture that element D is part of a young X-Y chromosome pair. Stable fly element D is also enriched for a unique set of transposable elements (e.g., *Vingi* and *Dada*) that are not enriched on any other element (Olafson *et al.*, 2019), which is also expected for an evolving sex chromosome (Ellison and Bachtrog, 2013; Steinemann and Steinemann, 2005). The stable fly X and Y chromosomes also likely both contain element F, which probably fused to element D, because element F genes have elevated male heterozygosity (**Fig 2B**).

The stable fly male-determining locus was previously mapped to chromosome 1 (Willis *et al.*, 1981). We therefore conclude that stable fly chromosome 1 corresponds to Muller elements D and F. The stable fly Y chromosome carries a male-determining locus, but we do not know the nature of this gene. The house fly male-determining gene (*Mdmd*) was not found in the stable fly genome or any other fly relatives (Sharma *et al.*, 2017). We also searched for the male-determining gene from tephritid flies (*MoY*; Meccariello *et al.*, 2019) in the stable fly genome using BLAST (Altschul *et al.*, 1990), and we failed to find anything resembling the *MoY* protein sequence. Therefore, either stable fly has an independently derived new male-determiner or it has retained an ancestral male-determining locus that was replaced in house fly.

### The horn fly sex chromosomes consist of elements A and F

We hypothesize that horn fly also has a young X-Y chromosome pair. The horn fly genome was sequenced using DNA extracted from a mixed sample of males and females (Konganti *et al.*, 2018), which prevents us from using heterozygosity in the genome sequencing reads to identify the horn fly sex chromosomes. However, there is available RNA-seq data from horn fly testis and ovary, which we used to identify the horn fly sex chromosomes using the same approach as we did in stable fly (Supplementary tables S2 and S5). In horn fly, there is elevated relative male heterozygosity in genes assigned to element A (*P* =0.024 in a Mann-Whitney test comparing element A with elements B–E) and element F (*P* =0.0059 in a Mann-Whitney test comparing element F with elements B–E), but none of the other major elements (**Fig 2C**). There is also a higher fraction of element A genes that only have heterozygous SNPs in males and not in females (385*/*889 = 43.3%) when compared to genes on the other major elements (1571*/*4163 = 37.7%; *z*=3.09, *P* =0.001). In addition, of the 44 horn fly genes assigned to element F, 11 have heterozygous SNPs. Of those 11 genes, 8 are only heterozygous in males, and the remaining 3 have more heterozygous SNPs in males than females. It is highly unlikely that >50% of heterogyzous SNPs would only be observed in males for all 11 of the horn fly element F genes if heterozygosity were equal in males and females (*z*=3.32, *P* =0.0005). There is also a higher fraction of element F genes that only have heterozygous SNPs in males (8*/*11 = 72.7%) when compared to genes on the autosomes (*z*=2.39, *P* =0.008). We therefore conclude that the horn fly X and Y chromosomes are likely both composed of elements A and F, and the sex chromosomes arose through a fusion of those two elements.

As with stable fly, we cannot examine heterozygosity along the full length of any individual Muller elements in horn fly because we lack a chromosome-scale assembly, and scaffold order along each chromosome has not been determined. Also, as in stable fly, male heterozygosity is, on average, higher than female heterozygosity across the entire horn fly genome (**Fig 2C**). Unlike stable fly, we cannot test for a more fragmented assembly of the horn fly sex chromosomes because the entire horn fly genome assembly is fragmented. The scaffold N50 of the horn fly assembly is 23 Kb (Konganti *et al.*, 2018), and the vast majority (4,112*/*4,778 = 86%) of horn fly scaffolds assigned to Muller elements have only 1 annotated gene. We also searched for both the house fly and tephritid male-determining genes in the horn fly genome using BLAST (Altschul *et al.*, 1990; Meccariello *et al.*, 2019; Sharma *et al.*, 2017), and we failed to find either.

### Sex-biased gene expression on the sex chromosomes

Genes that are expressed at different levels between females and males are said to have “sex-biased” expression (Parsch and Ellegren, 2013). Genes with male-biased (female-biased) expression are often under- (over-) represented on fly X chromosomes as a result of the haploid dose of the X chromosome in males or sex-specific selection pressures that prevent (favor) the evolution of male-biased (female-biased) expression on the X (Meiklejohn and Presgraves, 2012; Meisel *et al.*, 2012; Parisi *et al.*, 2003; Rice, 1984; Sturgill *et al.*, 2007). We cannot perform statistical tests for differentially expressed genes between males and females using available data from either stable fly or horn fly because only one replicate RNA-seq sample was collected for each tissue type from each sex for each species.

However, we can calculate the relative expression of genes in males and females 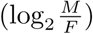 for each tissue type (Supplementary tables S6–S11), which differs between the X and autosomes in many flies (Vicoso and Bachtrog, 2015).

We compared the distributions of 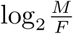 for genes assigned to each Muller element in stable fly, horn fly, and the closely related house fly (**Fig 4**). Genes on stable fly element D (part of the X and Y chromosomes) have significantly lower 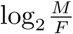 in reproductive tissues than genes on the other major elements (*P* = 0.0032 in a Mann-Whitney test). The slight but significantly reduced 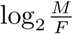 on element D is consistent with the “demasculinization” or “feminization” of the X chromosome observed in reproductive tissues of other flies (Meisel *et al.*, 2012; Parisi *et al.*, 2003; Sturgill *et al.*, 2007; Vicoso and Bachtrog, 2013, 2015). In contrast, genes on horn fly element F have higher 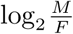 in gonad than genes on the autosomal elements (*P* =10^−4^ in a Mann-Whitney test), suggesting a “masculinization” of the ancestral X chromosome. The small number of annotated element F genes in stable fly likely limit our ability to detect significant masculinization of stable fly element F (**Fig 4**). There is no significant difference in 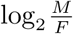 between genes on horn fly element A (the new portion of the sex chromosomes) and the autosomes. Element F is also part of a young sex chromosome pair in house fly (Meisel *et al.*, 2017). There is no evidence for a difference in 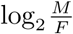 between genes on element F and the autosomes in house fly. The minimal evidence for demasculinization or feminization of the muscid X chromosomes is consistent with the sex chromosomes being diploid in both males and females in all three species. Similar expression in males and females is also consistent with young sex chromosomes that have not yet had time to accumulate sexually antagonistic alleles that could lead to sex-biased expression.

**FIG. 4.**
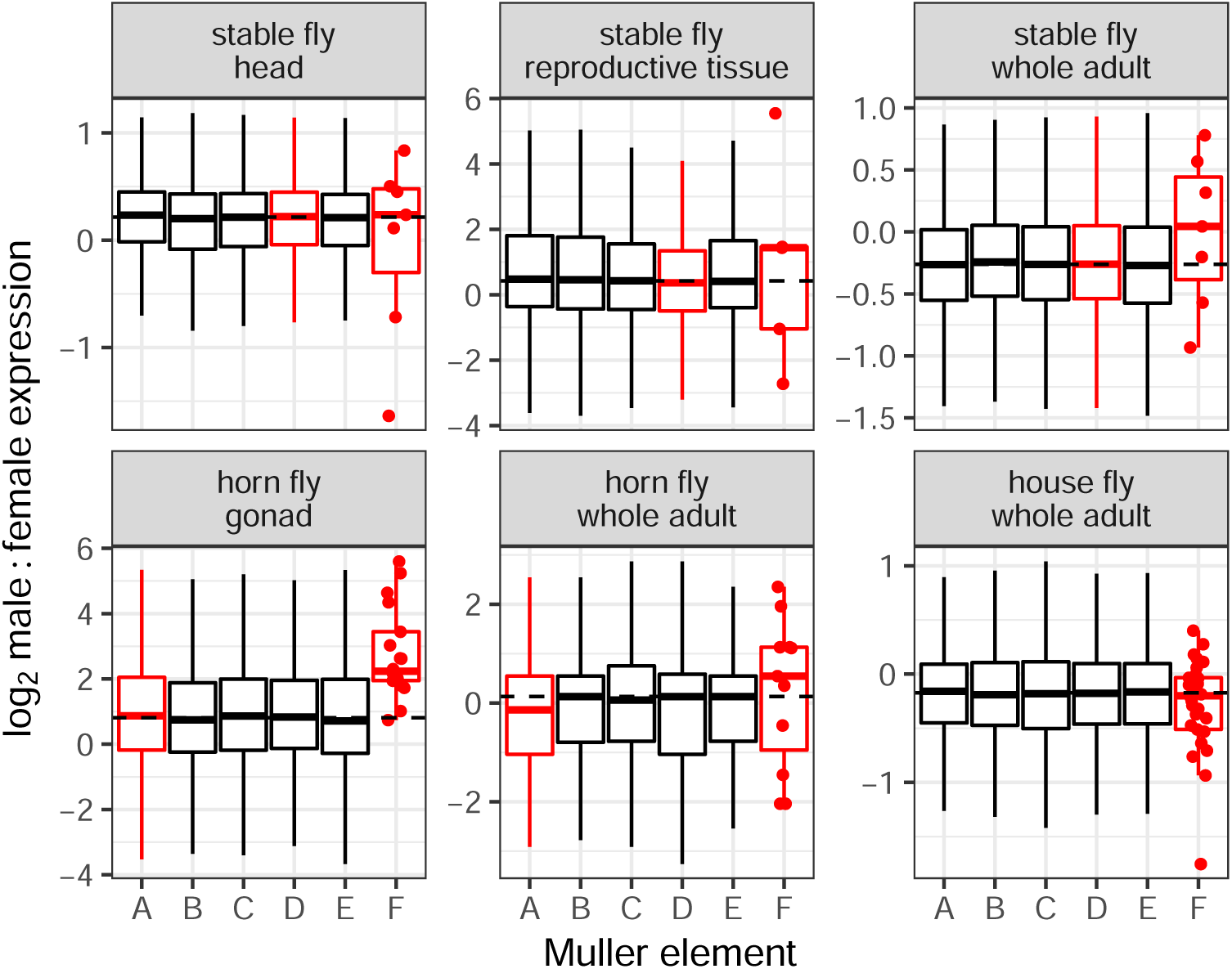
Sex-biased expression across Muller elements. The distributions of log_2_ male:female expression (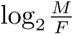) for genes on each Muller element in stable fly, horn fly, and house fly are shown with boxplots. Expression data are from either whole adult (all species), gonad (horn fly only), reproductive tissues (stable fly only), or head (stable fly only). Each data point used to generate the boxplots corresponds to an individual gene. Dots show the expression levels of individual genes on element F. Dashed lines indicate the genome-wide average for all genes. Outliers were omitted from the boxplots. Inferred sex-linked elements are drawn in red.

## Discussion

Stable fly and horn fly have derived karyotypes in which the ancestral X chromosome (Muller element F) and Y chromosome are not visible (Avancini and Weinzierl, 1994; Boyes *et al.*, 1964; Joslyn *et al.*, 1979; LaChance, 1964; Parise-Maltempi and Avancini, 2007). We show, based on elevated male heterozygosity, that the X and Y chromosomes of stable fly both contain elements D and F (**Fig 2A-B**). The reduced assembly quality of element D is further evidence that it is sex-linked in stable fly (**Fig 3**). We also present evidence that the X and Y chromosomes of horn fly contain elements A and F (**Fig 2C**). Elevated male heterozygosity is a hallmark of a young and undifferentiated sex chromosome pair (Vicoso and Bachtrog, 2015), suggesting that stable fly and horn fly have independently and recently derived nascent sex chromosomes. In addition, house fly also has multiple young proto-Y chromosomes (Meisel *et al.*, 2017). The minimal feminization/masculinization of gene expression on the sex chromosomes across these three species is consistent with their recent origins (**Fig 4**). House fly, stable fly, and horn fly diverged ∼27 My ago (Ding *et al.*, 2015), but the sex chromosomes in each species may have arisen more recently given the minimal X-Y sequence divergence in all three species.

Our evidence for the sex-linkage of element D in stable fly is strong because there is a consistent signal from absolute male heterozygosity (**Fig 2A**), relative male heterozygosity (**Fig 2B**), and assembly quality (**Fig 3**). In contrast, the only evidence for sex-linkage of element A in horn fly is elevated relative male heterozygosity (**Fig 2C**). However, there is support from other work that suggests elevated relative male heterozygosity is a reliable indicator of an undifferentiated X-Y pair (Meisel *et al.*, 2017; Toups *et al.*, 2019; Veltsos *et al.*, 2019; Vicoso and Bachtrog, 2015; Wright *et al.*, 2017; Yoshida *et al.*, 2014). We are therefore confident that element A is indeed sex-linked in horn fly. Future work could be done to further evaluate the sex-linkage of horn fly elements A and F, as well as stable fly elements D and F.

Curiously, we observe a pattern consistent with masculinization of the ancestral X chromosome (element F) in stable fly and horn fly (**Fig 4**), although only significant in horn fly due to a small sample size of stable fly element F genes. This is surprising because element F genes trend toward female-biased expression both in flies with the ancestral karyotype (X-linked element F) and in *Drosophila* where element F has reverted to an autosome (Vicoso and Bachtrog, 2013). The masculinization of element F in stably fly and horn fly suggests that a Y-linked copy of element F may have accumulated alleles that increase male expression (Zhou and Bachtrog, 2012). Alternatively, element F could be hyper-expressed in stable fly and horn fly males because it is both diploid and transcription is up-regulated by an ancestral dosage compensation system. Dosage compensation in a closely related blow fly, which has the ancestral fly karyotype (i.e., only element F is X-linked), is regulated by an RNA-binding protein that increases the transcriptional output of element F genes in hemizygous males (Davis *et al.*, 2018; Linger *et al.*, 2015). Stable fly and horn fly could have elevated element F expression in males because those genes are both up-regulated and diploid. Additional work is necessary to test these hypotheses.

We propose three different scenarios that could have given rise to the stable fly and horn fly cryptic sex chromosomes (**Fig 5**). The order of most events in each scenario is arbitrary, and it is not necessary for the sex chromosomes of stable fly and horn fly to have arisen by the same scenario. All three scenarios assume that the MRCA of muscid flies had a karyotype with five euchromatic autosomes (elements A–E) and a heterochromatic sex chromosome pair (where element F is the X chromosome) because this is the ancestral karyotype of Brachycera (Vicoso and Bachtrog, 2013, 2015) and is still conserved in some Muscidae (**Fig 1**). We additionally assume that the Y chromosome of the MRCA of Muscidae carried a male-determining locus because that is the most common mechanism of sex determination in closely and distantly related families (Bopp *et al.*, 2014; Meccariello *et al.*, 2019; Scott *et al.*, 2014b; Willhoeft and Franz, 1996).

**FIG. 5.**
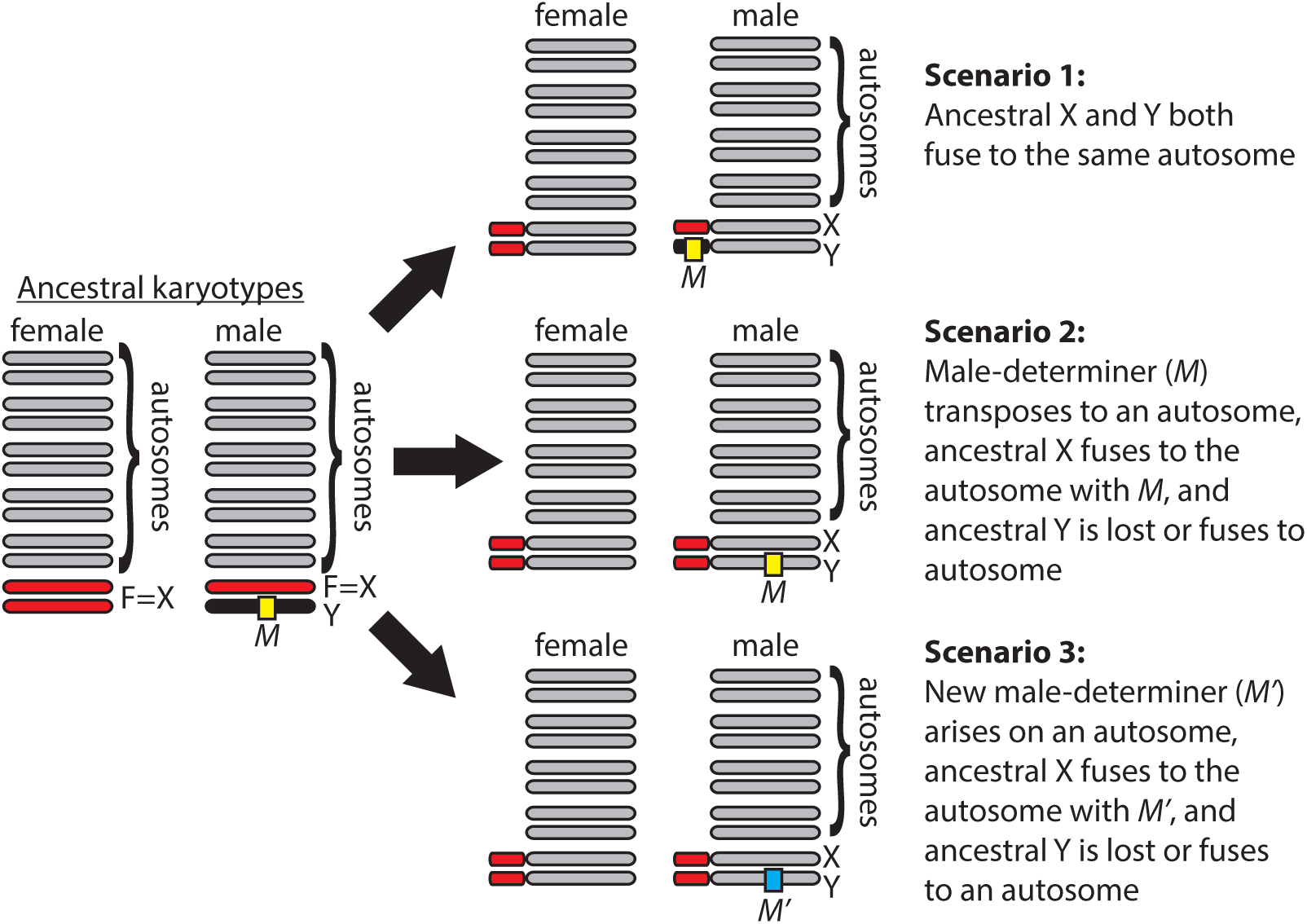
Potential scenarios that could give rise to muscid cryptic sex chromosomes. Ancestral autosomes (gray), X chromosome (red), and Y chromosome (black) are shown in both male and female karyotypes. Possible derived male and female genotypes are shown for three potential scenarios that could give rise to the cryptic sex chromosomes in stable fly and horn fly.

In the first scenario we hypothesize that both the X and Y chromosomes of the MRCA of Muscidae fused to the same ancestral autosome (**Fig 5**). These X-autosome and Y-autosome fusions would convert one copy of the ancestral autosome into a neo-X chromosome and the other copy into a neo-Y chromosome. Concurrent fusions between the X and Y to the same autosomal element may seem unlikely, but it has been observed in *Drosophila* and birds (Flores *et al.*, 2008; Pala *et al.*, 2011). Element F genes have elevated heterozygosity in both stable fly and horn fly males (**Fig 2**), suggesting that males carry two copies of element F that have the same gene content and are only slightly differentiated at the sequence level. Therefore, this scenario requires that the ancestral X and Y of Muscidae was undifferentiated, with the Y chromosome essentially just a copy of element F that carries a male-determining locus. The X and Y chromosomes of house fly are very similar in gene content (Meisel *et al.*, 2017), but we do not know if this is the ancestral state of Muscidae or a derived condition in house fly.

In the second scenario, we hypothesize that the ancestral male-determining gene transposed from the ancestral Y chromosome to an autosome (**Fig 5**). This transposition event would convert the autosome into a proto-Y chromosome, and its homolog would be a proto-X chromosome. A transposing male-determining gene (*Mdmd*) was identified in house fly (Sharma *et al.*, 2017), demonstrating the feasibility of this scenario. We also hypothesize that the ancestral X chromosome (element F) fused to the same autosome containing the male-determining locus in this scenario. We acknowledge that element F fusing to the same chromosome that carries the male-determining gene would be a remarkable coincidence. Moreover, element F must have fused to both a copy of the autosome with the male-determiner (the proto-Y) and a copy of the autosome without a male-determiner (the proto-X). This is because males must have two differentiated copies of element F (one X-linked and the other Y-linked) in order to explain the elevated male heterozygosity of element F genes in both stable fly and horn fly (**Fig 2**). The order of the F-autosome fusion and the transposition of the male-determiner is arbitrary in this model.

In the last step of the second scenario, the ancestral Y chromosome either fused to one of the autosomes or it was lost from the genome. Such a transposition of Y chromosome genes to an autosome happened following the creation of a neo-X chromosome in *Drosophila pseudoobscura* (Carvalho and Clark, 2005; Larracuente *et al.*, 2010). A Y-autosome fusion is possible in this scenario if the Y chromosome lost its male-determining activity, possibly via a pseudogenizing mutation. Alternatively, the ancestral Y chromosome could have fused to the element carrying the transposed male-determiner (the proto-Y), which would allow the ancestral Y to retain the male-determiner without creating an independently segregating second Y chromosome. If the ancestral Y chromosome was lost, we hypothesize that the ancestral Y did not contain any essential genes other than the male-determiner. Some *Drosophila* species have Y chromosomes that lack essential genes (Voelker and Kojima, 1971), demonstrating that it is feasible for a fly Y chromosome to not be essential for male viability or fertility. Moreover, the genetic differentiation of X and Y chromosomes in both blow fly and flesh fly could be explained by a lack of essential genes on their Y chromosomes other than the male-determiner (Linger *et al.*, 2015; Vicoso and Bachtrog, 2013, 2015).

The third scenario differs from the second in that instead of the ancestral male-determiner transposing to an autosome, a new male-determiner arises on one of the autosomes (**Fig 5**). The new male-determiner would convert the autosome into a proto-Y chromosome, and its homolog would be a proto-X. The male-determining *Mdmd* gene in house fly arose from a highly conserved splicing factor that was duplicated after the divergence between house fly and stable fly (Sharma *et al.*, 2017), demonstrating that new male-determining genes can arise within Muscidae. As in the second scenario, element F would have fused to the same chromosome carrying the new male-determiner (i.e., the proto-Y), and it must have also fused to the homologous proto-X to produce the elevated male heterozygosity we observe in element F genes (**Fig 2**). Once again, like the second scenario, fusion of element F to the same chromosome that carries the male-determining gene would be a remarkable coincidence. In addition, the ancestral Y was either lost or fused to an autosome, as in the second scenario.

In all three scenarios, invasion (and fixation) of the fusion between element F and an autosome may be favored if one copy (more likely a Y-autosome fusion, but also possibly an X-autosome fusion) confers a sex-specific fitness benefit (Charlesworth and Charlesworth, 1980; Matsumoto and Kitano, 2016). The effect of sex-specific selection on the invasion of Y-autosome fusion will be greater if there is no recombination between the neo-X and neo-Y (Charlesworth and Charlesworth, 1980). One way for recombination to be suppressed is if there is no recombination in males, as is the case in *Drosophila* and many other flies (Gethmann, 1988). There have been no tests for male recombination in either stable fly or horn fly, but there is some evidence for male recombination in the closely related house fly (Feldmeyer *et al.*, 2010). Testing for male recombination in other muscid flies will be important for evaluating which selective forces could be responsible for the new sex-linked elements in Muscidae.

Each of the three scenarios makes predictions about the sex chromosomes of the MRCA of Muscidae, and testing those predictions is necessary in order to evaluate the hypotheses. For example, if all muscid species with the ancestral karyotype have undifferentiated X and Y chromosomes (as in house fly), then we would infer that the MRCA had undifferentiated sex chromosomes, supporting scenario 1. Alternatively, if extant muscids with the ancestral karyotype have differentiated X and Y chromosomes, then we could infer that the MRCA had differentiated X and Y chromosomes. This would support scenarios 2 and 3.

In addition, identifying the male-determining genes across Muscidae would allow us to test if the same gene is used for male-determination across species or if new male-determiners have arisen in species other than the house fly (Sharma *et al.*, 2017). If the same male-determiner is used across species, then scenarios 1 and 2 would be supported. If new male-determiners arose in species with derived karyotypes, then scenario 3 would be supported. Therefore, by characterizing the sex chromosomes and male-determining genes across muscid flies, we can distinguish between all three scenarios. Distinguishing between the scenarios would provide valuable insights into the factors that promote sex chromosome evolution in muscid flies, which would serve as an informative model for understanding why rates of sex chromosome evolution differ across taxa. For example, is a high rate of sex chromosome evolution in Muscidae promoted by ancestrally undifferentiated X and Y chromosomes, a gene-poor ancestral Y, transposing male-determiners, a high rate of new male-determiners, or some combination of multiple factors? Testing these hypotheses should motivate future work in this system.

In summary, we have identified independently derived young sex chromosomes in stable fly and horn fly (**Fig 2**), which are different from the proto-sex chromosomes of house fly (Meisel *et al.*, 2017; Sharma *et al.*, 2017). Therefore, there are at least three independently derived young sex chromosome systems in Muscidae, and probably more based on the derived karyotypes distributed across the family (**Fig 1**). In addition, we present three possible scenarios for the origins of the stable fly and horn fly sex chromosomes (**Fig 5**). Each scenario makes specific predictions about the male-determining genes and sex chromosomes in the MRCA of Muscidae and in other muscid species. Notably, the scenarios include two important factors that could allow for faster rates of sex chromosome evolution in some taxa (e.g., Muscidae) than in closely related taxa (e.g., blow flies and flesh flies). First, the ancestral X and Y chromosomes of Muscidae could have been undifferentiated, in contrast to the differentiated blow fly and flesh fly sex chromosomes (Davis *et al.*, 2018; Linger *et al.*, 2015; Vicoso and Bachtrog, 2013, 2015). Undifferentiated sex chromosomes could allow for the formation of new sex chromosomes (Dufresnes *et al.*, 2015; Stöck *et al.*, 2011, 2013). Second, there could be a high rate of new or transposing male-determining genes across Muscidae, as is the case in house fly (Sharma *et al.*, 2017). These new or transposable male-determining genes could allow allow for a faster rate of sex chromosome turnover. Testing for undifferentiated sex chromosomes, new male-determining genes, and transposing male-determiners in Muscidae is therefore a promising approach to assess the relative importance of these factors in permitting or promoting frequent and rapid sex chromosome turnover.

## Acknowledgments

Some of the computational analyses reported were performed on the Maxwell Cluster that is part of the University of Houston Research Computing Data Core. We thank T. J. Raszick, S.-H. Sze, C. J. Coates, and A. M. Tarone for sharing results on repeat content in the stable fly genome.

